# Stability of phoneme-related potentials across testing sessions and stimulus presentation conditions

**DOI:** 10.64898/2026.05.26.727924

**Authors:** Zhe-chen Guo, Kailyn A. McFarlane, Jacie R. McHaney, Ishika Choksi, Mollee Feeney, Lauren Preston, Bharath Chandrasekaran

**Affiliations:** Roxelyn and Richard Pepper Department of Communication Sciences and Disorders, Northwestern University

**Keywords:** phoneme-related potentials, electroencephalography, speech perception, continuous speech, clinical translation

## Abstract

**Objectives:** Objective and ecologically valid measures of speech processing can complement conventional audiologic assessments. Phoneme-related potentials (PRPs), derived by averaging listeners’ electroencephalography (EEG) responses time-locked to phonemes in continuous speech, have emerged as a promising approach for capturing cortical processing of speech in naturalistic listening conditions. Importantly, PRPs reveal speech perception challenges even when conventional audiograms are clinically normal, positioning them as a promising neural marker for suprathreshold listening difficulties that standard audiometry often misses. As a critical step toward clinical translation, this study examined the extent to which PRP-derived measures remain stable across real-world contexts relevant to clinical implementation, including monaural versus binaural presentation, stimulus intensity level, and repeated testing sessions. The study also assessed cortical tracking of lower-level speech acoustics to determine whether the PRP findings could be attributed to acoustic processing.

**Design:** EEG was recorded from 18 young adults with normal hearing as they listened to audiobook speech presented monaurally or binaurally at 60 or 75 dB across two sessions separated by approximately one week. Neural differentiation of phoneme manner-of-articulation classes (vowels, nasals/approximants, fricatives, and stops) in PRPs was quantified using two measures: an *F*-statistic reflecting between-manner relative to within-manner variability, and classification accuracy from a machine-learning model trained to predict manner class from PRPs. Temporal response function modeling assessed neural tracking of continuous acoustic envelope and onset features of the audiobook speech.

**Results:** Neither PRP-derived measure of manner differentiation showed significant effects of session, presentation modality, intensity level, or their interactions. Intraclass correlation analyses further indicated moderate-to-good reliability across all three factors. In contrast, neural tracking of the acoustic envelope and acoustic onsets was stronger under binaural than monaural presentation, with binaural presentation eliciting more pronounced cortical responses to the envelope.

**Conclusions:** PRP-derived measures remained relatively stable across modest procedural variations that are common in clinical testing contexts, positioning PRPs as a potent objective index of naturalistic speech processing. This stability may reflect cortical processing of abstract, linguistically relevant speech categories and suggest that PRPs provide complementary information beyond audiologic assessments of peripheral auditory functions and EEG measures that primarily capture lower-level acoustic processing.

## Introduction

A key mission of hearing healthcare revolves around facilitating spoken language communication. A multitude of clinical populations experience significant speech perception challenges. Yet, the gold standard of hearing assessment—pure-tone audiometry—often fails to capture the extent of speech perception abilities (Killion et al., 2004). Current clinical assessments of speech perception vary in the linguistic and cognitive strategies they engage (DiNino et al., 2022), often lack ecological validity and a neurobiological basis, and require a behavioral response from the patient, posing additional challenges for certain clinical populations. Thus, there is a clear need for an objective, ecologically valid assessment of speech processing that can be used to complement standard approaches.

Advances in electrophysiological methods have increasingly moved the field toward more ecologically valid assessments of speech processing (see Hamilton & Huth, 2020, for a review). Rather than relying solely on simple tones or isolated speech sounds that have been the backbone of prior electrophysiological approaches, newer approaches emphasize neural responses to continuous, naturalistic speech. Among these approaches, neural tracking of continuous speech has emerged as a powerful framework for assessing the extent to which the brain encodes dynamic speech signals. Polonenko and Maddox (2021) developed a method for capturing subcortical responses to continuous speech by pairing re-synthesized “peaky” speech with deconvolution analysis of the auditory brainstem response. At the cortical level, studies have shown that neural responses recorded via electroencephalography (EEG) closely track the amplitude envelope of continuous speech (Aiken & Picton, 2008; Di Liberto et al., 2015; Ding & Simon, 2014). This cortical tracking is commonly quantified by fitting a temporal response function (TRF; Brodbeck et al., 2023; Crosse et al., 2016) model to describe the linear mapping relationship between the envelope and neural activity. Because the envelope contains acoustic cues to speech features including syllables and word boundaries (Oganian & Chang, 2019), TRF-based measures of envelope tracking have been used to provide an index of cortical speech processing and to examine how it is altered by factors including aging (Decruy et al., 2019; Gillis et al., 2023; McHaney et al., 2021) and hearing loss (Fuglsang et al., 2020).

Beyond tracking the continuous speech envelope, an emerging approach with translational potential is to examine neural responses to phonemes, the building blocks of words (e.g., *cat* consists of the phonemes K, AE, and T). Intracranial studies in human patients have revealed that speech perception is subserved by distributed neural ensembles within the superior temporal gyrus (STG) that are selectively tuned to the acoustic-phonetic properties of speech sounds. This selectivity within temporal lobe populations show that phoneme representations are an emergent property of distributed activity across neural ensembles tuned to specific speech features (Gnanateja et al., 2025; Leonard et al., 2015, 2024; Leonard & Chang, 2014; Mesgarani et al., 2014; Mesgarani & Chang, 2012). Grounded in this neurobiological framework, recent efforts have examined phoneme discrimination using non-invasive EEG. Scalp-recorded EEG is collected while participants listen to continuous speech stimuli, such as an audiobook. Event-related potentials time-locked to instances of each phoneme are extracted and then averaged to derive “phoneme-related potentials” (PRPs; Khalighinejad et al., 2017; Figure 1A). PRPs to individual phonemes or to phoneme classes have been used as a proxy of neural differentiation of speech sounds, a key precursor to behavioral speech perception. Recent work by Guo et al. (2025) suggests that PRPs reveal subtle, subclinical speech perceptual challenges in middle age not detected by audiometry. Specifically, Guo et al. found that despite clinically normal hearing thresholds, middle-aged adults showed poorer words-in-noise perception than younger adults and a concomitant reduction in neural differentiation of speech sound categories indexed by PRPs to continuous speech. Furthermore, these less distinct phoneme representations in middle-age could not be entirely explained by peripheral auditory measures or cortical tracking of low-level acoustic features such as the speech envelope. Other studies have similarly reported less distinct PRPs in clinical populations including cochlear implant users (Aldag & Nogueira, 2024) and under degraded listening conditions (Jeon & Woo, 2023). Together, these findings highlight PRPs as a promising addition to clinical assessments, with the potential to capture perceptual difficulties with meaningful speech categories under naturalistic listening conditions in ways that traditional audiological test batteries may not.

**Figure 1.**
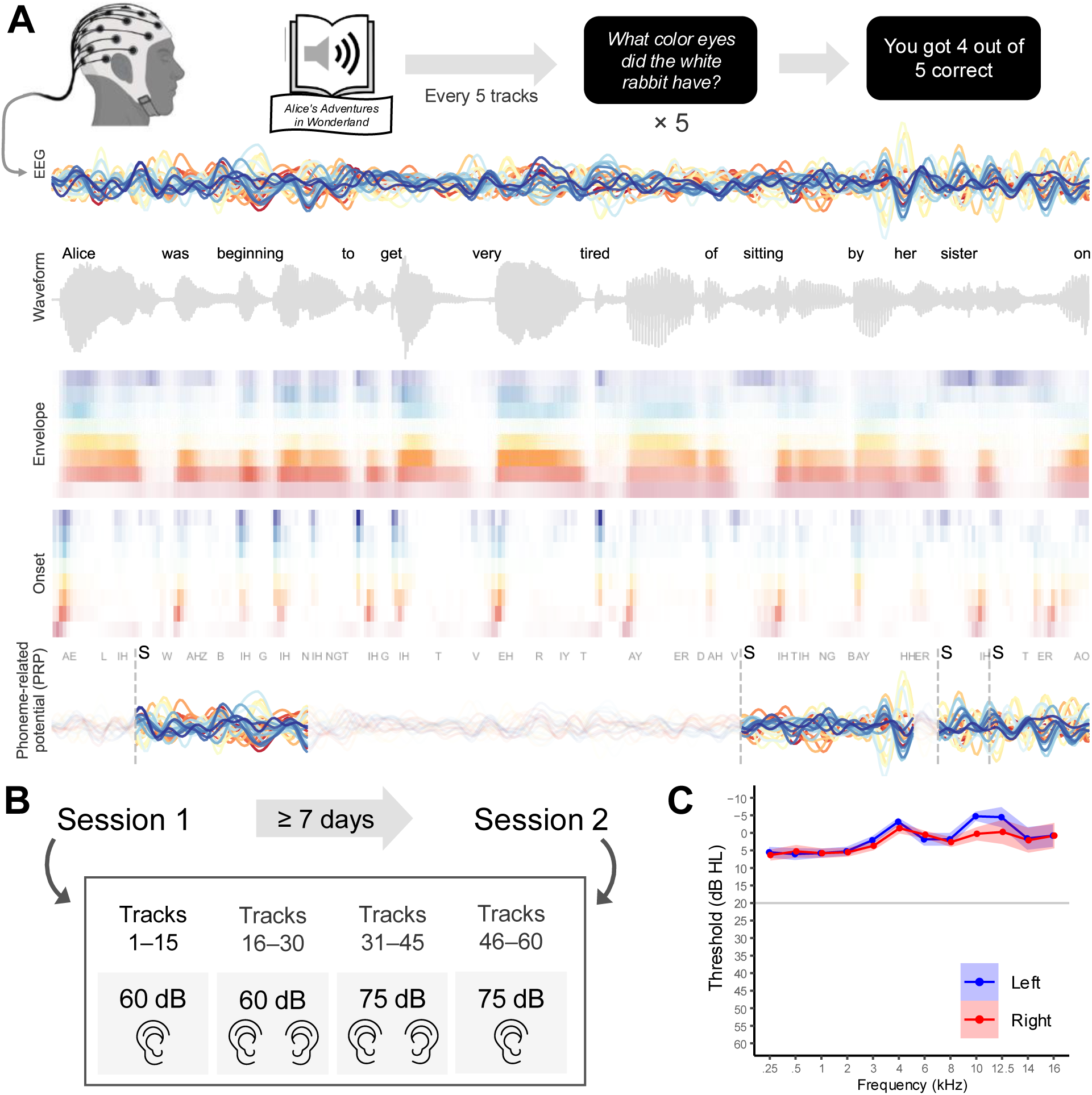
Continuous speech listening experiment. (A) EEG was recorded while participants listened to a public-domain audiobook of Alice’s Adventures in Wonderland (Carroll, 1865) in quiet. The audiobook was segmented into 60 tracks of approximately 60 s each. After every five tracks, participants answered five comprehension questions assessing their understanding of the narrative. The panel illustrates a representative segment of processed EEG data from one participant alongside the time-aligned speech waveform and the corresponding acoustic envelope and onset features across eight frequency bands. Phoneme-related potentials were extracted by averaging EEG responses time-locked to all instances of a given phoneme (e.g., S) within a 0–500 ms window following phoneme onset. (B) Participants listened to 15 consecutive tracks under each combination of presentation mode (monaural, binaural) and intensity level (60 dB, 75 dB), with tracks presented in chronological order to preserve the storyline. Approximately one week later, participants returned for a second session, during which the same procedure was repeated. (C) Average bilaterial hearing thresholds between 0.25–16 kHz of the participants.

However, translating PRPs to clinical use requires a better understanding of common factors that can impact the robustness of a measure in the clinical context. In the present study, we examine three factors with direct relevance to clinical implementation: monaural versus binaural presentation, stimulus intensity, and test-retest reliability.

While many EEG studies present continuous speech stimuli binaurally, the majority of audiologic tests are administered monaurally to obtain ear-specific diagnostics. Demonstrating that PRPs remain a robust neural measure of speech category separability under monaural presentation would facilitate integration into routine clinical workflows and broaden their applicability to populations for whom ear-specific assessment is especially valuable. These include individuals with unilateral hearing loss, single-sided deafness, as well as cochlear implants. Monaural presentation also allows aided-versus-unaided comparisons in hearing aid users, who often suffer from significant asymmetries between ears. At the same time, binaural listening improves speech intelligibility through binaural summation and redundant acoustic input (Bronkhorst & Plomp, 1989; Kokkinakis & Pak, 2014; Marks, 1978), potentially also enhancing PRPs. Reduced speech category separability under monaural presentation would then suggest that binaural listening contributes meaningfully to neural differentiation of speech categories. Thus, we ask whether PRPs can effectively capture speech category separability under monaural presentation, or whether binaural presentation is necessary to preserve this distinction.

Another key consideration is stimulus intensity. Using moderate levels ensures comfort and feasibility for patients with hyperacusis or tinnitus, pediatric listeners, and individuals with reduced sound tolerance. To obtain more robust PRPs, higher intensities may be used as they are known to increase the strength of cortical auditory evoked potentials to tone stimuli (Beagley & Knight, 1967; Billings et al., 2007). However, some evidence suggests that cortical responses may instead be reduced at higher intensities, revealing suprathreshold auditory deficits due to saturation, central gain compensation, or reduced access to high-threshold auditory-nerve fibers. Animal models show that even when hearing thresholds are normal or have recovered, noise-induced hidden hearing loss alters neural discriminability of speech sounds in a level-dependent manner, reducing it at a higher presentation level while enhancing it at a moderate level (e.g., 75 vs. 60 dB; Monaghan et al., 2020). Together, these findings suggest that the neural coding patterns of speech sounds can vary as a function of suprathreshold intensity. Assessing the stability of PRPs across intensity levels thus help determine whether they can be flexibly adjusted to meet the needs of the listener, and whether they may provide a framework protocol for using different intensities to identify suprathreshold auditory dysfunction.

Finally, test-retest reliability is essential for any measure to be used for diagnosis, longitudinal monitoring, or evaluation of treatment outcomes. Ideally, PRPs can be collected using standardized continuous speech stimuli (e.g., audiobook passages) that could be used across repeated testing sessions. If repeated testing does involve the same speech materials, it is important to determine whether increased familiarity with the content influences PRPs independently of auditory function, given that it could modulate engagement, attention, arousal, prediction, or listening effort (Brown et al., 2020; Rommers & Federmeier, 2018; Tremblay et al., 2010; Vanden Bosch der Nederlanden et al., 2022). Demonstrating that PRP-derived measures are also stable across sessions would strengthen confidence that these measures reflect relatively trait-like aspects of speech encoding rather than state fluctuations.

To this end, we recorded EEG from young adults with normal hearing thresholds while they listened to continuous speech in quiet, presented across four conditions formed by crossing two presentation modalities (monaural and binaural) and two intensity levels (60 dB and 75 dB). Participants returned to complete the same four conditions in a second session approximately one week later. We focused on quantifying the neural differentiation of natural speech sound classes defined by manner of articulation that are robustly represented in PRPs (Aldag & Nogueira, 2024; Guo et al., 2025; Khalighinejad et al., 2017): vowels, nasals and approximants, fricatives, and stops. This classification has practical clinical relevance because audiological speech-assessment materials, leverage broad phonetic contrasts to evaluate access to speech cues that differ in spectral and temporal properties (Scollie et al., 2012). Accordingly, manner-based PRP differentiation may provide a clinically interpretable index of how well the auditory system preserves the acoustic-phonetic structure common to everyday speech perception. We examined whether PRP-based measures of manner differentiation remain stable across presentation modalities, stimulus intensity levels, and sessions.

We also used TRF modeling to assess the extent to which these factors influenced cortical tracking of envelope-based acoustic features. This measure has also been examined as a potential translational marker of continuous speech processing and shown to remain stable across testing sessions separated by at least one week (Dial & Gnanateja, 2025; Panela et al., 2024). However, it remains unclear whether envelope tracking is similarly stable across monaural versus binaural presentation and across different stimulus intensity levels. Importantly, assessing cortical tracking of speech acoustics will help determine whether any differences or lack of differences in PRP-derived measures across conditions could be attributed to lower-level acoustic processing or instead reflect speech category processing beyond acoustics.

## Methods

### Participants

Eighteen native English speakers (11 females; age range: 19–26 years, *M* = 21.8, *SD* = 2.19) participated in the multi-session study. To be included, participants were required to have normal hearing (see Audiological assessments). Four additional individuals were recruited but excluded for not completing all four conditions in either session (*N* = 2), not returning for the second session of the experiment (*N* = 1), or for scoring outside normal limits on speech in noise testing (*N* = 1). All participants provided written informed consent and received monetary compensation for their time. All experimental procedures were approved by the Northwestern University Institutional Review Board (STU00222157).

### Audiological assessments

All participants were assessed for normal auditory function through a battery of audiological and electrophysiological measures. Prior to testing, otoscopy was performed to confirm participants’ ear canals were unobstructed. Air conduction audiometry was administered in a double-walled, sound-treated room using the Intelligent Hearing Systems (IHS; Miami, FL, USA) SmartAud software module (version 5.56.00) through the Duet platform. Stimuli were presented through foam-tip earbuds attached to ER-2 transducers (Etymotic Research, Elk Grove Village, IL, USA). Bilateral thresholds across 0.25–16 kHz were obtained via the modified Hughson-Westlake procedure. All participants included in the final analysis had bilateral thresholds ≤ 25 dB hearing level (HL) between 0.25–8 kHz (Figure 1C).

Word recognition in noise was tested using the Words in Noise test (WIN; Wilson et al., 2003), where participants repeated target words fixed at a level of 65 dB SPL against a multi-talker babble masker that systematically increased in level, decreasing the signal-to-noise ratio SNR) from 24 to 0 dB in 4-dB steps. Participants completed two test lists. The SNR loss in dB was calculated for each participant, which reflects the optimal SNR level necessary to achieve 50% accuracy. All participants included in the final analysis scored within the established normal limits (< 6 dB SNR; (Wilson et al., 2003)). The mean WIN SNR loss was 3.00 dB SNR with a standard deviation of 1.42.

Auditory brainstem responses (ABR) evoked by a 100 μs broadband click were collected from both ears. The stimulus was presented in alternating polarity at a rate of 19.3/s at 86 dB nHL through a gold-foil TIPtrode attached to a shielded ER-3C transducer. Responses were collected through the IHS Smart EP module (version 5.54.23) using a vertical electrode montage (active Cz, reference ipsilateral ear canal, Fpz ground), amplified at a 10^5^ gain and band-pass filtered between 30–3000 Hz. Per ear, two repeatable waveforms were collected and summed to obtain a grand average response (2048 sweeps total). Wave I and V absolute peak latencies were identified by an experienced electrophysiologist via visual inspection and marked using visual overlay cursors in the SmartEP software. All participants included in the final analysis had absolute and interpeak latencies within normal limits (Sanfins et al., 2022; Table 1).

**Table 1.** Auditory brainstem response absolute and interpeak latencies.

| Ear | Left Ear |  |  | Right Ear |  |  |
| --- | --- | --- | --- | --- | --- | --- |
| Wave | I | V | I-V<br>Interpeak | I | V | I-V<br>Interpeak |
| Mean (SD) | 1.55<br>(0.16) | 5.55<br>(0.25) | 4.00<br>(0.21) | 1.56<br>(0.12) | 5.54<br>(0.19) | 3.98<br>(0.14) |
| Range (min,<br>max) | 1.30, 2.04 | 5.20, 6.18 | 3.72, 4.65 | 1.33, 1.83 | 5.22, 5.85 | 3.73, 4.25 |

### Continuous speech listening experiment

The stimuli were continuous speech excerpts from the public-domain audiobook *Alice’s Adventures in Wonderland* (Carroll, 1865) narrated by a male speaker of American English and recorded at a sampling rate of 22.05 kHz. The audiobook has been widely used in prior research investigating the cortical processing of naturalistic speech (Gnanateja et al., 2025; Guo et al., 2025; Llanos et al., 2021; McHaney et al., 2021, 2025; Xie et al., 2023). Long pauses in the stimuli were truncated to a maximum of 500 ms and the entire audiobook was divided into 60 tracks of ∼60 s (range: 59–65 s). Word and phoneme segmentations were obtained using the Montreal Forced Aligner (McAuliffe et al., 2017).

The 60 tracks were arranged into four sets of 15 consecutive tracks while preserving their original chronological order to maintain the storyline (Figure 1B). Each set was presented in one of four condition combinations resulting from the two intensity levels (60 dB, 75 dB) and two presentation modalities (monaural, binaural). For example, a participant might hear tracks 1–15 monaurally at 60 dB, tracks 16–30 binaurally at 75 dB, tracks 31–45 monaurally at 75 dB, and tracks 46–60 binaurally at 60 dB. There were four unique orders of the four stimulus presentation conditions, and participants were assigned to one of these orders. Every five tracks, participants answered five four-alternative forced-choice comprehension questions about the story they had heard. Following the first session, participants returned at least one week later (mean 7.67 days, range: 7–13 days) for a second session to listen to the same tracks in the same condition order. This design yielded eight unique conditions (two sessions × two presentation modalities × two intensity levels). All stimuli were presented in quiet through ER-3C insert earphones (Etymotic Research, Elk Grove Village, IL) to both ears in the binaural condition or to the right ear only in the monaural condition.

### Acoustic envelope and onsets

Continuous acoustic envelope and onsets were extracted from the audiobook speech (Figure 1A) for subsequent neural tracking analyses (see Section 2.6.3). These acoustic features were generated using gammatone filters implemented in the Python Eelbrain package (Brodbeck et al., 2023), which simulated frequency response characteristics in the cochlea. The audiobook stimuli were passed through a 256-channel gammatone filterbank spanning 0.02–5 kHz to produce a high-resolution, 256-band spectrogram. The spectrograms were then downsampled to 1 kHz for computational efficiency, log-transformed, and collapsed into eight logarithmically spaced frequency bands to yield a multiband gammatone envelope. Because cortical responses are sensitive to acoustic onsets or rapid envelope changes (Daube et al., 2019; Hamilton et al., 2018; Yi et al., 2019), we also computed representations of acoustic onsets using an auditory edge-detection model (Fishbach et al., 2001). The resulting 256-band onset spectrograms were similarly summed into eight logarithmically spaced bands to obtain multiband acoustic onset features. Both the envelope and onset representations were further downsampled to 128 Hz to match the sampling rate of the cleaned EEG data.

### EEG data acquisition and preprocessing

EEG was recorded using a BioSemi ActiveTwo system (version 3.2; BioSemi B.V., Amsterdam, Netherlands) with 16-bit resolution at a sampling rate of 16,384 Hz. Participants wore a BioSemi headcap with fixed electrode positions, with the cap centered by aligning the Cz electrode midway between the nasion-inion axis and between the two ears. Cap size was selected based on each participant’s head circumference (nasion to inion). Recordings were obtained from 32 Ag/AgCl electrodes arranged according to the International 10–20 system (Klem et al., 1999). Electrode offsets were maintained below 25 mV for all channels. Two mastoid electrodes (left and right) and two periocular electrodes (below and lateral to the left eye) were used for offline referencing and ocular monitoring. EEG data were preprocessed in Python using the MNE software package (Gramfort, 2013). The data were first referenced to the common average, downsampled to 128 Hz for computational efficiency, and band-pass filtered from 1 to 15 Hz using minimum-phase, causal windowed-sinc FIR filters. Ocular artifacts were attenuated using regression. Independent component analysis (ICA) was subsequently applied using a face ICA method with 800 iterations to identify and remove components associated with muscle activity and eye movement before reconstructing the cleaned EEG.

### Data analyses

#### Analyses of phoneme-related potentials

For each phoneme, PRPs were obtained by averaging EEG signals time-locked to the onsets of all instances of that phoneme within a 0–500 ms window. This procedure was applied at each electrode for each participant and condition. Because phoneme occurrence frequencies vary substantially in naturalistic speech, we included only phonemes that accounted for at least 1% of all phoneme instances and excluded the schwa AH, which was the most frequent phoneme and accounted for approximately 10% of instances. This selection yielded a set of 31 phonemes, which can be grouped into four classes based on manner of articulation: 12 vowels (AA, AO, AW, AY, AE, EH, ER, EY, IH, IY, OW, UW), six nasals and approximants (M, N, NG, L, R, W), seven fricatives (S, SH, F, HH, Z, DH, V), and six stops (P, T, K, B, D, G). Consequently, the PRP dataset for each participant and condition consisted of 31 500-ms EEG responses across all 32 electrodes, representing typical neural responses to these phonemes.

Previous work has demonstrated that PRPs primarily reflect distinctions in manner of articulation, and this neural differentiation is reduced due to hearing loss (Aldag & Nogueira, 2024) and aging (Guo et al., 2025). Building on this evidence, we first examined whether session, presentation modality, and intensity level influenced manner-class separability of PRPs. Separability was quantified using the *F*-statistic, which is defined as the ratio of between-class variability to within-class variability (Patel et al., 1976). *F*-statistics were computed from PRP amplitude values for the 31 phonemes at each electrode and time point, separately for each participant and condition. To identify spatiotemporal patterns of significant effects, we conducted a mass-univariate ANOVA using Eelbrain (Brodbeck et al., 2023) on the resulting *F*-statistics of manner separability. Statistical significance was evaluated using a cluster-based permutation approach which controls for multiple comparisons by finding meaningful “clusters,” or contiguous electrodes showing the same effect (Maris & Oostenveld, 2007). *F*-values from the ANOVA test at uncorrected level of *p* ≤ 0.05 were used as the cluster forming threshold. Significant clusters (*p* ≤ 0.05) were then determined by comparing its cluster-mass statistic against a null distribution generated from 10,000 random permutations of the data. The maximum *F*-value within each cluster (*F*_max_) was reported as an effect size estimate (Brodbeck et al., 2018).

In addition to the mass-univariate tests conducted across the full spatiotemporal arrays, we performed a targeted analysis focused on three temporal landmarks across eight frontocentral electrodes (Fz, F3, F4, FC1, FC2, Cz, C3, and C4). These landmarks occur approximately at 50, 120, and 230 ms and are referred to as R1, R2, and R3, respectively, and prior work has shown that the time points exhibit distinct *F*-statistic peaks indicating maximal manner separability (Khalighinejad et al., 2017). The frontocentral electrodes were selected as the region of interest (ROI) because they are sensitive to manner-of-articulation contrasts in PRPs and show robust cortical responses to auditory stimuli (Khalighinejad et al., 2017; Panela et al., 2024; Picton et al., 2003). For each participant and condition, *F*-statistics were averaged across 20-ms windows centered on the R1, R2, and R3 peaks defined from the present data and then averaged across the ROI. These values were analyzed using linear mixed-effects models implemented with the lme4 package in R (Bates et al., 2015; R Core Team, 2023). Session, presentation modality, intensity level, and their interactions were included as fixed effects. Random effects included by-participant random intercepts and the maximal random-slopes structure for the fixed factors permitted by the model and data. Statistical significance of fixed effects was assessed using Type-III ANOVA with Satterthwaite’s approximation.

We also employed a supervised machine-learning classification approach (Aldag & Nogueira, 2024; Guo et al., 2025) to assess the extent to which manner classes could be discriminated based on PRPs. PRPs from the frontocentral ROI electrodes were pooled across participants, and mean amplitudes were extracted within 20-ms windows centered on the R1, R2, and R3 peaks. These three values served as features for classification. Classification was performed using a support vector machine (SVM) with a radial basis function kernel and evaluated using 10-fold cross-validation. At each iteration, 10% of the data were held out as the test set. Because the four manner classes differed in the number of constituent phonemes, phonemes from overrepresented classes were randomly subsampled to create a balanced dataset with an equal number of phonemes per class. From the remaining data, another 10% was reserved as a validation set, and the rest was used to train the classifier. Hyperparameter tuning was conducted over a range of values for the regularization parameter *C* = {0.01, 0.1, 1, 10, 100}, and the model yielding the highest validation accuracy was selected to predict manner-class labels for the test set. This procedure was repeated until all data had served as test sets and was performed separately for each of the eight conditions. Because phoneme balancing and data partitioning involved random sampling, the entire classification pipeline was repeated 30 times using different random seeds. Classification accuracy was then averaged across repetitions, electrodes, and phonemes to yield a single accuracy value for each participant in each condition. These accuracy values were analyzed using a linear mixed-effects model to assess the effects of session, presentation modality, intensity level, and their interactions.

The manner classification model yielded not only accuracy rates but also confusion patterns. We thus further examined whether the patterns of classification confusion differed as a function of session, presentation modality, or intensity level using a permutation analysis. For each participant and condition, a four-by-four confusion matrix was constructed and normalized such that each row (corresponding to the true manner class) summed to one. To assess the effect of a given factor (e.g., session), confusion matrices of each participant were first averaged separately for the two levels of that factor (e.g., session 1 and session 2), collapsing across the levels of the remaining two factors. From each of the resulting averaged matrices, cell values were extracted and flattened into a one-dimensional vector. The Spearman correlation distance (1 − *ρ*) between the two vectors was computed. The mean distance across participants served as the observed difference in classification patterns between the two levels of the factor of interest. A null distribution of mean distances was generated using 10,000 permutations to evaluate statistical significance. In each permutation, the labels of the factor of interest were randomly shuffled within each participant prior to averaging the confusion matrices and computing the correlation distance. The permutation *p*-value was then calculated as the proportion of permuted distances that were greater than or equal to the observed mean distance.

#### Reliability of PRP-derived measures

We also directly quantified the reliability of the above PRP-derived measures across sessions, presentation modalities, and intensity levels using intraclass correlation coefficients (ICC). We focused on: 1) average *F*-statistic of manner separability at R1, R2, and R3 peaks in the *F*-statistic time course; 2) manner classification accuracy. To evaluate reliability across a given factor, the values of each measure from each participant were averaged separately for the two levels of that factor collapsing across the levels of the remaining two factors. ICC was calculated using single measurement, absolute agreement, two-way mixed-effects models (ICC(2,1); Koo & Li, 2016; Shrout & Fleiss, 1979), treating individual participants as observations. Reliability was interpreted as excellent for ICC >0.9, good for ICC between 0.75 and 0.9, moderate for ICC between 0.5 and 0.75, and poor for ICC < 0.5 (Koo & Li, 2016).

#### Neural tracking of continuous acoustic features

Finally, given that auditory cortical activity tracks the spectrotemporal dynamics of continuous speech such as the amplitude envelope (Aiken & Picton, 2008; Di Liberto et al., 2015; Lalor & Foxe, 2010), we assessed participants’ neural tracking of the continuous acoustic envelope and acoustic onsets. This analysis was included to determine whether the observed patterns in PRP-derived measures could be more parsimoniously explained by differences (or lack thereof) in the processing of lower-level acoustic dynamics. To address this question, we fitted an encoding a multivariate TRF model (Crosse et al., 2016) to predict the EEG signal (*t*) from the multiband acoustic envelope and onsets. Formally, the model is given by:

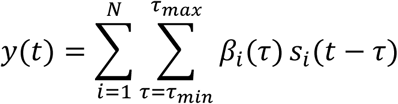

where *s_i_*(*t*) denotes the *i*-th stimulus feature (with *N* = 16 corresponding to envelope and onset features across the eight bands), and *β_i_*(*τ*) is the temporal response function associated with feature *i* at time lag *τ*, representing the influence or “weights” of each elementary event in this feature over time. The lag window was defined from *τ_min_* = −100 ms to *τ_max_* = 500 ms. The EEG response is thus modeled as the sum of the convolutions between each stimulus feature and its corresponding TRF. The TRF model was estimated separately for each electrode, participant, and experimental condition using the boosting algorithm in the Eelbrain package (Brodbeck et al., 2023) with five-fold cross-validation and an *ℓ*1 error norm. Prediction accuracy of the estimated model was quantified as the Pearson’s correlation (*r*) between the predicted and observed EEG signals, with a higher correlation indicating more robust neural tracking of the acoustic features.

To assess the effects of session, presentation modality, intensity level, and their interactions on neural tracking, we performed a mass-univariate ANOVA on the prediction-accuracy topographies, following the same procedure used for the PRP analysis. We also conducted ROI analysis focusing on the frontocentral ROI electrodes by fitting a mixed-effects model to prediction accuracy values averaged across the ROI for each participant and condition. In addition, we examined whether the estimated TRF curves varied as a function of session, presentation modality, and intensity level. For each acoustic predictor type, envelope or onset, TRFs were first averaged across the eight frequency bands to yield a single TRF at each electrode for each participant and condition. Mass-univariate ANOVAs were then used to identify spatiotemporal clusters showing significant main effects or interactions.

## Results

### PRP-derived measures

Figure 2 shows the grand average PRP to each phoneme and each manner of articulation across all four conditions and two sessions. *F*-statistics quantifying neural separability of the four manner classes over time (Figure 3A) exhibited three prominent peaks at 55 ms, 125 ms, and 211 ms, referred to as R1, R2, and R3, respectively. The latencies of these peaks closely matched those reported in prior studies (Khalighinejad et al., 2017; R1: 70 ms, R2: 130 ms, R3: 200 ms). Mass-univariate ANOVA tests conducted on the entire spatiotemporal *F*-statistic arrays found no clusters showing significant effects of session, presentation, intensity level, or their interactions (session: *F*_max_ = 4.424, *p* = 0.521; intensity: *F*_max_ = 5.421, *p* = 0.393; presentation: *F*_max_ = 1.713, *p* = 1.000; session × intensity: *F*_max_ = 4.675, *p* = 0.480; session × presentation: *F*_max_ = 7.676, *p* = 0.274; intensity × presentation: *F*_max_ = 1.783, *p* = 1.000; session × intensity × presentation: *F*_max_ = 4.538, *p* = 0.504; Figure 3C). Focusing on average *F*-statistics around R1, R2, and R3 in the ROI (Figure 3B), mixed-effects analysis similarly provided no evidence for any significant main effect or interaction (session: *F*(1, 17) = 0.013, *p* = 0.909; intensity: *F*(1, 102) = 0.062, *p* = 0.804; presentation: *F*(1, 102) = 0.275, *p* = 0.601; session × intensity: *F*(1, 102) = 0.003, *p* = 0.956; session × presentation: *F*(1, 102) = 0.976, *p* = 0.326; intensity × presentation: *F*(1, 102) = 1.868, *p* = 0.175; session × intensity × presentation: *F*(1, 102) = 0.264, *p* = 0.609).

**Figure 2.**
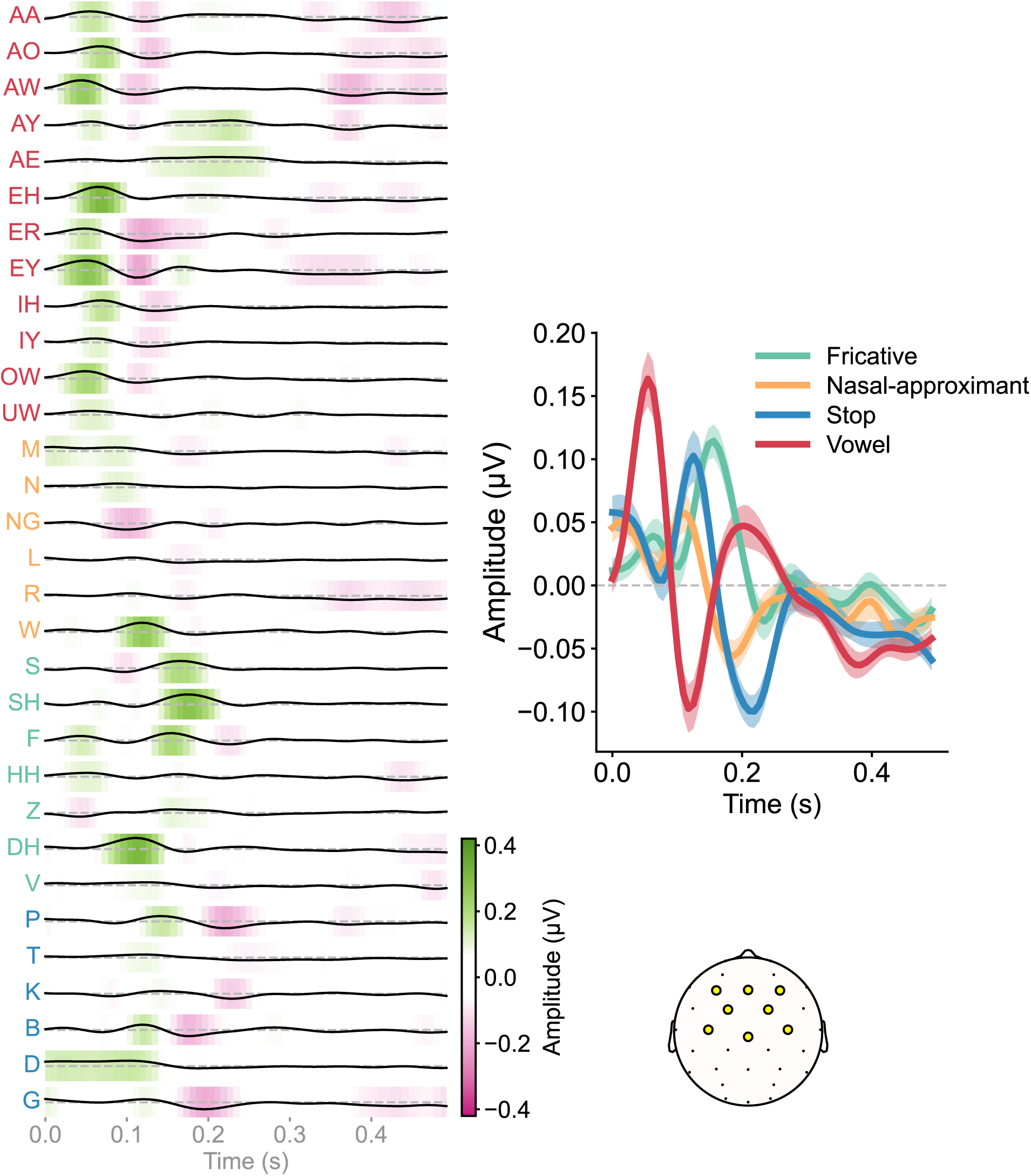
Phoneme-related potentials (PRPs) to each phoneme and each manner of articulation, averaged across all participants and conditions and across the eight frontocentral region-of-interest (ROI) electrodes (yellow circles in the scalp topography). Phoneme PRP amplitudes are color-coded to aid visualization. Shaded regions indicate ±1 standard error of the mean.

**Figure 3.**
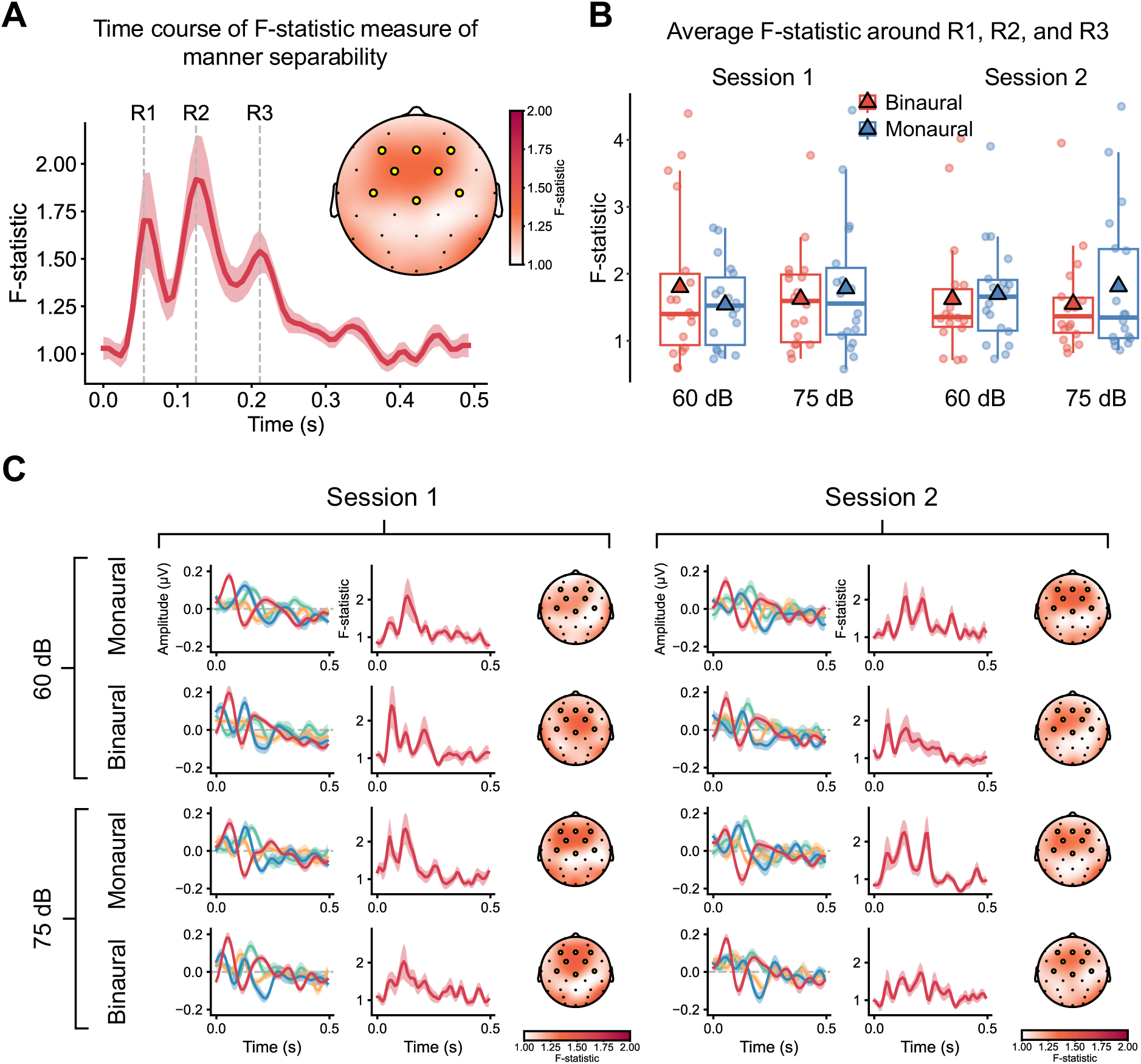
*F*-statistic measures of phoneme-related potentials (PRPs). (A) Time course of the F-statistic averaged across participants, conditions, and the eight frontocentral region-of-interest (ROI) electrodes (yellow circles in the scalp topography). The topographic map (inset) shows the scalp distribution of the *F*-statistic averaged across the PRP time window. The first three peaks in the *F*-statistic time course are labeled R1, R2, and R3. (B) Boxplots of *F*-statistic values averaged within 20-ms windows centered on R1, R2, and R3 across the ROI electrodes. Circles denote individual participants, and triangles indicate the mean for each condition. (C) PRPs grouped by manner of articulation, corresponding *F*-statistic time courses, and time-averaged *F*-statistic scalp topographies for each combination of session, presentation modality, and intensity level. Shaded regions indicate ±1 standard error of the mean.

Figure 4A presents the accuracy of the classifier trained to predict manners from PRPs along with the classification confusion matrices averaged across all conditions (Figure 4B) and divided by session, presentation modality, and intensity level (Figure 4C). Mixed-effects analysis revealed no significant effects or interactions (session: *F*(1, 17.002) = 2.392, *p* = 0.140; intensity: *F*(1, 84.997) = 0.088, *p* = 0.768; presentation: *F*(1, 17.001) = 0.217, *p* = 0.647; session × intensity: *F*(1, 84.997) = 1.122, *p* = 0.293; session × presentation: *F*(1, 84.997) = 0.519, *p* = 0.473; intensity × presentation: *F*(1, 84.997) = 0.889, *p* = 0.349; session × intensity × presentation: *F*(1, 84.997) < 0.001, *p* = 0.989). Permutation tests indicated that the classification confusion patterns were not significantly affected by session (permutation *p* **=** 0.105), presentation modality (permutation *p* **=** 0.074), or intensity level (permutation *p* **=** 0.360).

**Figure 4.**
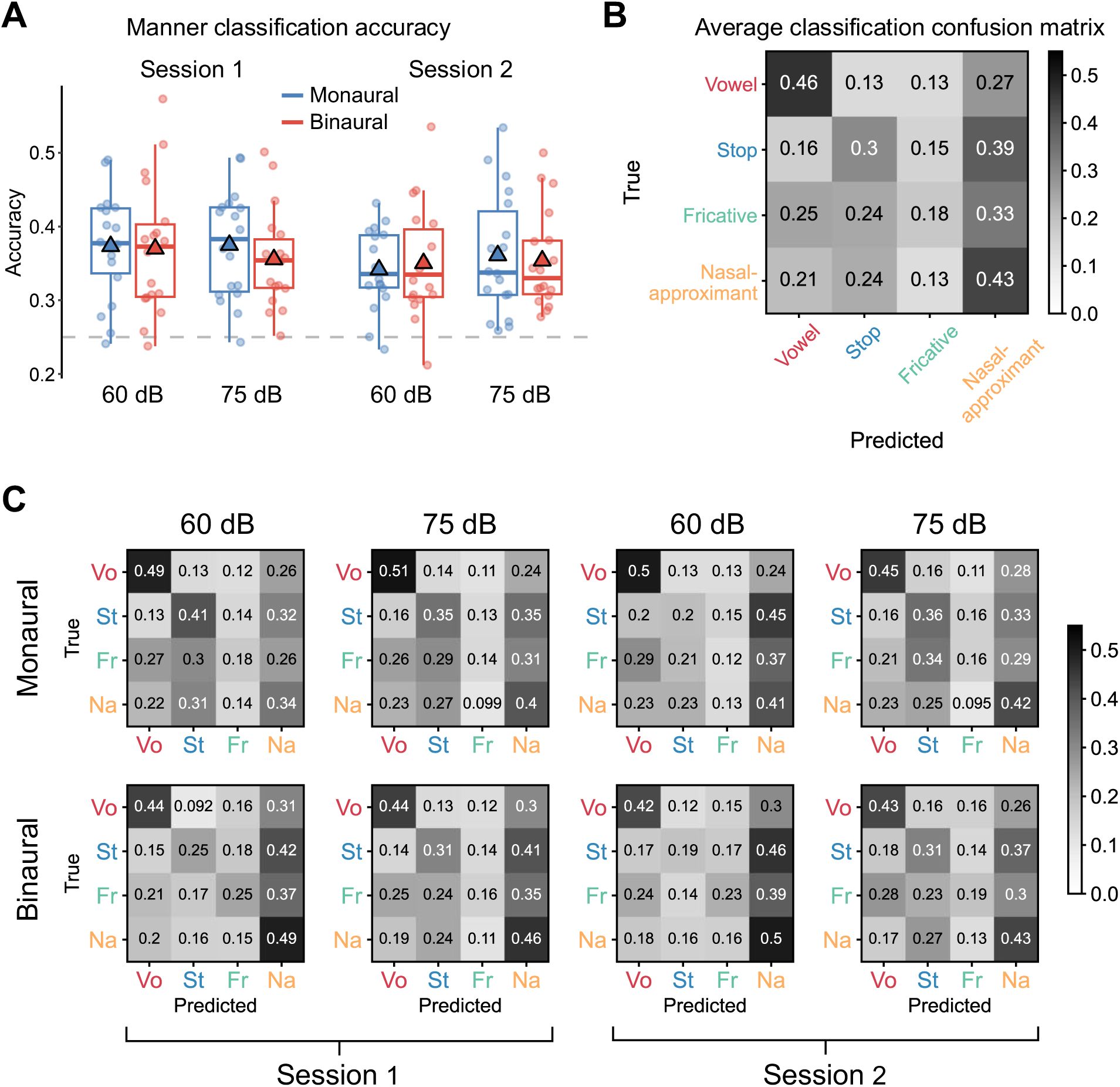
Phoneme-related potential (PRP) manner classification results. (A) Boxplots of classification accuracy from a machine-learning classifier trained to predict phoneme manner. Circles denote individual participants, and triangles indicate the mean for each condition. (B) Confusion matrix of classification results averaged across all conditions. (C) Confusion matrices shown separately for each condition. Values in each cell are normalized by the true class labels. *Vo*. = vowel; *St*. = stop; *Fr*. = fricative; *Na*. = nasal-approximant.

Figure 5 summarizes the ICC results of manner separability (average *F*-statistic around R1, R2, and R3 at the ROI electrodes) and manner classification accuracy. For manner separability, ICCs ranged from 0.640 (moderate) to 0.753 (good), with the highest reliability observed across presentation modalities. For manner classification accuracy, ICCs ranged from 0.652 (moderate) to 0.838 (good), with the highest reliability observed across intensity levels. All ICC estimates were statistically significant (all *p*s < 0.01). Overall, these results indicate moderate-to-good reliability for both PRP-derived measures (Koo & Li, 2016).

**Figure 5.**
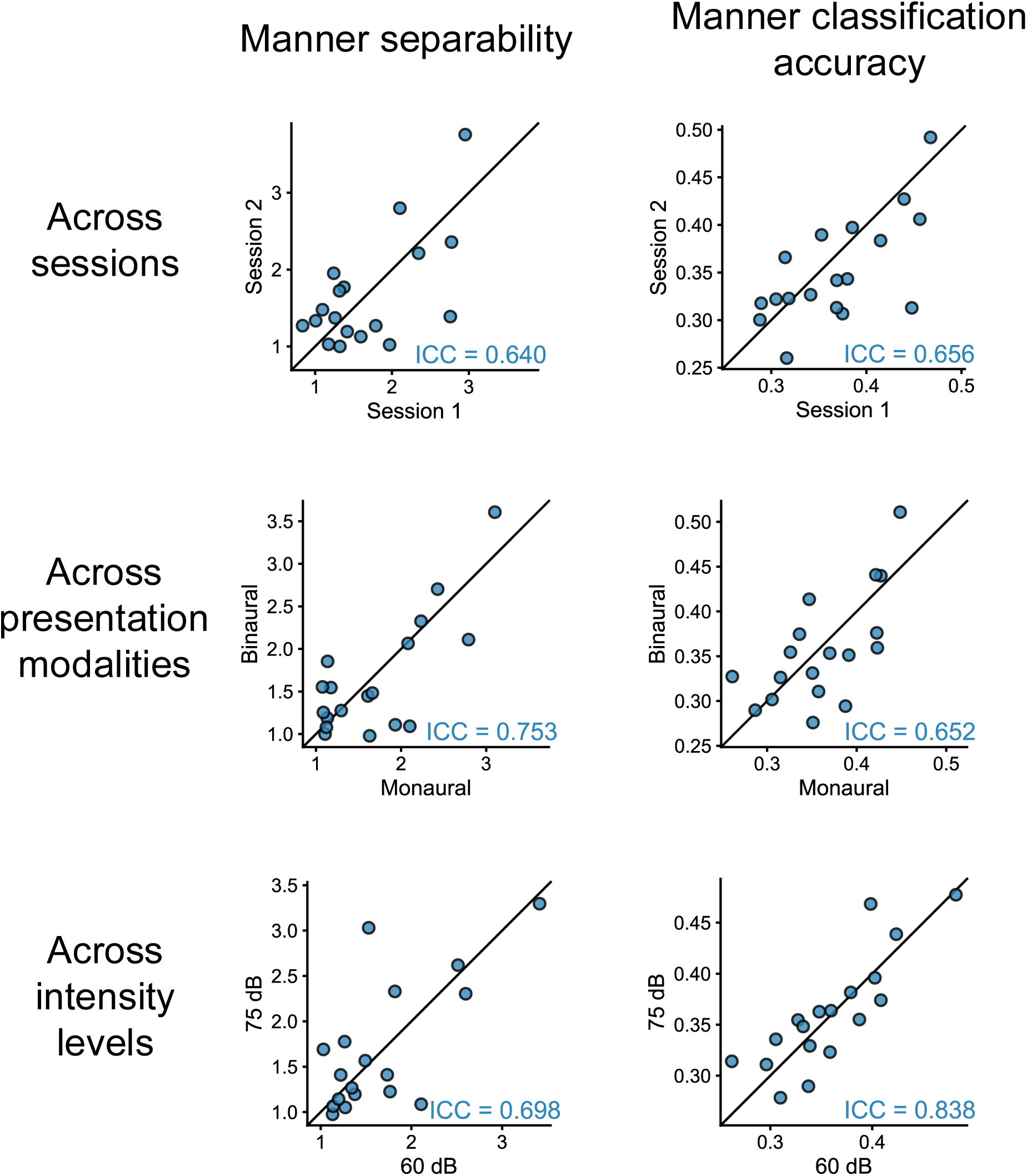
Reliability of measures derived from phoneme-related potentials across experimental factors. Scatterplots show manner separability and manner classification accuracy across sessions, presentation modes, and intensity levels for individual participants (circles), along with corresponding intraclass correlation coefficients (ICC).

### Neural tracking of acoustic envelope and onsets

Figure 6A shows the mean accuracy (Pearson’s correlation *r*) of the TRF model predicting EEG responses from the multiband acoustic envelope and onsets, divided by session, presentation modality, and intensity level. Mass-univariate ANOVA tests revealed a cluster where prediction accuracy was significantly higher in the binaural than monaural condition (*F*_max_ = 8.807, *p* = 0.049, 4 electrodes). No significant clusters were found for the remaining main effects or interactions (session: *F*_max_ = 6.210, *p* = 0.166; intensity: *F*_max_ = 2.076, *p* = 1.000; session × intensity: *F*_max_ = 2.586, *p* = 1.000; session × presentation: *F*_max_ = 2.128, *p* = 1.000; intensity × presentation: *F*_max_ = 2.436, *p* = 1.000; session × intensity × presentation: *F*_max_ = 2.252, *p* = 1.000). For prediction accuracy averaged across the frontocentral ROI (Figure 6B), linear mixed-effect modeling revealed a significant main effect of presentation modality (*F*(1, 17.001) = 11.161, *p* = 0.004), with higher accuracy under binaural than monaural presentation. A significant interaction between session and intensity level was also observed (*F*(1, 67.996) = 5.282, *p* = 0.025), indicating that the effect of intensity differed between sessions. Specifically, prediction accuracy at 75 dB was reduced relative to 60 dB in session 2 compared with session 1. No other main effects or interactions were significant (session: *F*(1, 17.001) = 3.357, *p* = 0.084; intensity: *F*(1, 17.003) = 0.472, *p* = 0.501; session × presentation: *F*(1, 67.996) = 0.023, *p* = 0.880; intensity × presentation: *F*(1, 67.996) = 0.745, *p* = 0.391; session × intensity × presentation: *F*(1, 67.996) = 0.996, *p* = 0.322).

**Figure 6.**
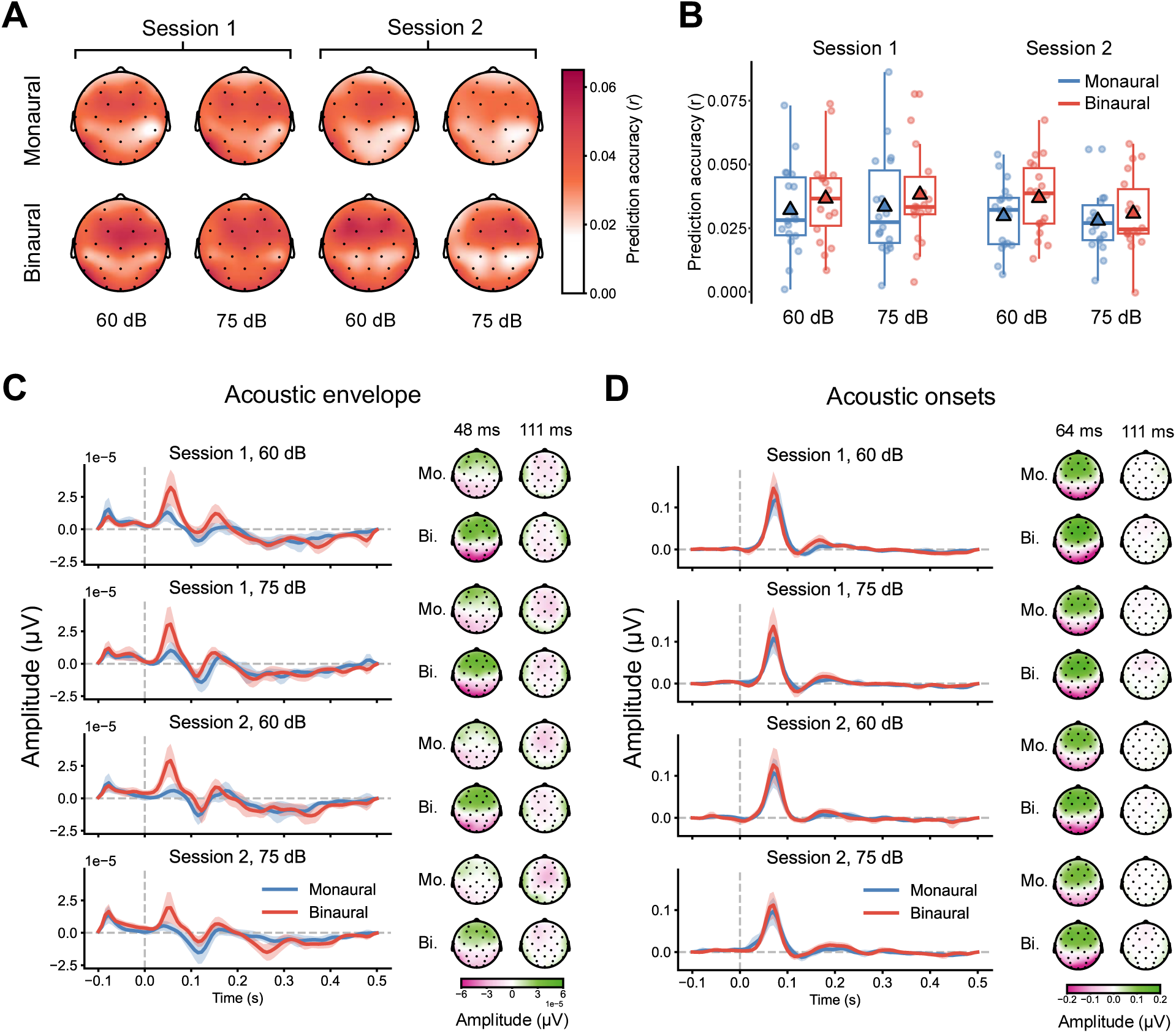
Neural tracking of continuous acoustic envelope and onset features. (A) Scalp topographies of prediction accuracy (r) from multivariate temporal response function (TRF) models predicting EEG responses from acoustic envelope and onset features across eight frequency bands, averaged across participants for each combination of session, presentation modality, and intensity level. (B) Boxplots of prediction accuracy averaged across the eight frontocentral electrodes defining the region of interest (ROI). Circles represent individual participants, and triangles indicate the mean for each condition. (C) Estimated TRFs for the acoustic envelope, averaged across frequency bands and across electrodes within the frontocentral ROI. Topographic maps to the right of the TRF waveforms depict scalp distributions of TRF amplitudes at the first positive peak (P1; 48 ms) and the subsequent negative peak (N1; 111 ms). *Mo*. = monaural; *Bi*. = binaural. (D) Corresponding TRF waveforms and P1 (64 ms) and N1 (111 ms) scalp topographies for acoustic onset features.

Figures 6C and 6D show the mean estimated TRFs, averaged across frequency bands, within the frontocentral ROI for the acoustic envelope and acoustic onset features, respectively. Mass-univariate ANOVA revealed that the spatiotemporal TRF profiles for the acoustic envelope were significantly modulated by presentation modality. Specifically, a significant cluster was identified for this effect in the time window from −0.022 to 0.181 s (*F*_max_ = 30.884, *p* < 0.001, 27 electrodes). For the acoustic onset TRFs, significant differences were also observed across presentation modalities, with one cluster extending from 0.119 to 0.236 s (*F*_max_ = 15.431, *p* = 0.013, 13 electrodes). In addition, TRF profiles for acoustic onsets varied as a function of intensity level in a separate cluster from −0.077 to 0.041 s (*F*_max_ = 13.706, *p* = 0.025, 13 electrodes). No significant clusters were found for other main effects or interactions in the TRF analyses of either acoustic envelope (session: *F*_max_ = 15.276, *p* = 0.466; intensity: *F*_max_ = 13.865, *p* = 0.507; session × intensity: *F*_max_ = 14.664, *p* = 0.131; session × presentation: *F*_max_ = 12.221, *p* = 0.523; intensity × presentation: *F*_max_ = 10.625, *p* = 0.895; session × intensity × presentation: *F*_max_ = 8.994, *p* = 0.670) or acoustic onset (session: *F*_max_ = 12.912, *p* = 0.318; session × intensity: *F*_max_ = 13.084, *p* = 0.259; session × presentation: *F*_max_ = 12.306, *p* = 0.839; intensity × presentation: *F*_max_ = 10.452, *p* = 0.704; session × intensity × presentation: *F*_max_ = 9.802, *p* = 0.798). These results indicated that neural tracking of continuous envelope-based speech features was modulated by the experimental factors particularly presentation modality, with binaural presentation resulting in higher prediction accuracy and more pronounced envelope TRFs than monaural presentation.

## Discussion

The larger goal of this work is to determine the extent to which PRPs can serve as reliable, clinically interpretable markers of neural processing of speech. Toward that goal, we examined the stability of PRP-derived measures across stimulus presentation conditions and sessions, contexts that have high clinical relevance. EEG was recorded from young, normal-hearing adults as they listened to continuous speech presented monaurally or binaurally at 60 or 75 dB, and the same procedure was repeated approximately one week later. PRP-derived measures of manner-of-articulation class separability were remarkably stable: neither the *F*-statistic measure nor machine-learning classification accuracy was significantly affected by session, presentation modality, intensity level, or their interactions. ICC analyses indicated moderate-to-good reliability for both these PRP-derived measures across all three factors. TRF analyses showed that cortical tracking of continuous multiband acoustic envelope and onset features was stronger under binaural than monaural presentation, with a significant session-by-intensity interaction indicating reduced difference between the two intensity levels in the second session. Together, these findings suggest that PRPs provide a reliable index of speech category processing in naturalistic listening, supporting their potential as an objective, ecologically valid complement to current clinical assessment approaches.

From a theoretical perspective, the stability of PRP-derived measures across presentation modalities, intensity levels, and repeated sessions suggests that PRPs capture a relatively invariant level of speech processing. Since speech discriminability is a crucial precursor to speech perception, PRPs may be well-suited for clinical deployment, where testing conditions often induce variability. The same factors influenced neural tracking of the continuous acoustic envelope and onsets. In particular, binaural presentation, likely due to summation, elicited stronger envelope tracking and more pronounced envelope TRFs than monaural presentation to the right ear. This difference aligns with the well-documented advantages of binaural listening in behavioral speech perception and the evidence for distinct auditory evoked-response profiles under binaural versus monaural stimulation (Avan et al., 2015; Balkenhol et al., 2020; Ching et al., 2004; Hawley et al., 2004). Therefore, although neural tracking of the speech envelope features are reliable across sessions (Dial & Gnanateja, 2025; Panela et al., 2024) and reveals acoustic processing differences across listener populations (e.g., younger and older adults; Decruy et al., 2019; Gillis et al., 2023), our results suggest that presentation modality would need to be factored in for clinical application.

Together, the findings indicate that continuous-speech EEG measures are not uniformly sensitive to the same processes and may reflect distinct underlying neurobiological mechanisms. Whereas envelope tracking captures broader sensory-attentional entrainment to the acoustic stream, PRPs likely index a more discrete and feature-specific stage of cortical speech representation that supports phonemic categorization. This dissociation is consistent with intracranial studies showing partially distinct functional and anatomical organization of acoustic and phonological representations within human auditory cortex, particularly along the STG. Direct cortical recordings indicate that posterior STG responds preferentially to acoustic onsets, while cortical sites selective for phonemes or phonetic features are more broadly distributed and intermixed across posterior and anterior lateral STG (Gnanateja et al., 2025; Hamilton et al., 2021; Yi et al., 2019). Moreover, cortical response patterns to phonemes are organized primarily according to manner-of-articulation distinctions (Mesgarani et al., 2014), consistent with the manner-level separability captured by PRPs. A similar dissociation has been reported for pitch-related prosodic features, such that some cortical sites preferentially encode absolute pitch associated with speaker identity, whereas others encode speaker-normalized relative pitch supporting linguistic intonational contrasts (Tang et al., 2017). Thus, measures derived from PRPs time-locked to discrete phoneme categories and neural tracking of the continuous acoustic stream may provide complementary, rather than redundant, information about speech processing.

The relative insensitivity of PRP-derived measures to modest procedural variation has important clinical implications. Because these measures remained stable across the factors examined here, they may be easier to standardize across audiology clinics, patient populations, and longitudinal follow-up visits than measures that are more sensitive to specific testing configurations. This flexibility may be valuable in several clinical scenarios. For example, the stability of PRPs under monaural presentation suggests that they could be extended to assessments in patients with unilateral hearing loss. For listeners with hyperacusis, presentation levels could be adjusted to improve comfort and feasibility without substantially affecting PRP-derived outcomes. In addition, because animal models of hidden hearing loss show level-dependent abnormalities in suprathreshold neural coding (Monaghan et al., 2020), the stability across the 60 dB and 75 dB conditions in our young normal-hearing sample may provide a normative benchmark. Deviations from this stable pattern in listeners with normal audiograms could indicate subtle suprathreshold dysfunction. The observed test-retest reliability further suggests that the PRP-derived measures are not strongly affected by short-term fluctuations in stimulus familiarity or listener state but can reveal changes that occur over longer timescales and affect speech sound representations, such as reduced neural differentiation of phoneme categories in middle age (Guo et al., 2025). This property makes the PRP approach useful for monitoring progression of central auditory decline, hearing intervention outcomes, rehabilitation, and auditory training. Together, the findings support the potential of PRPs as a scalable and ecologically valid marker of speech processing in naturalistic listening contexts.

For clinical adoption, the next key step is to establish a feasible recording setup. Most EEG studies of continuous speech perception rely on research-grade systems with 32 or 64 electrodes, which are often too time-consuming to set up and difficult to incorporate into routine clinical protocols. Additional barriers include the need for personnel training and the cost of specialized EEG equipment. However, translational success has been demonstrated for some EEG-based measures including the ABR. The ABR provides clinically relevant information about subcortical speech processing using a minimal montage of only a few electrodes placed at locations such as Cz and the mastoids (Polonenko & Maddox, 2021; Russo et al., 2004). For the PRP approach, electrodes over the frontocentral scalp may be sufficient to capture the most relevant information, as this region consistently shows robust neural separability of manner classes in PRPs as well as strong neural tracking of the speech envelope (Khalighinejad et al., 2017; Panela et al., 2024; Picton et al., 2003). Because our ROI analyses were based on only eight frontocentral electrodes, the present findings raise the possibility that an even sparser ABR-like montage—for example, with just the Cz and Fz electrodes referenced to mastoids—may be able to recover reliable PRP-based measures of speech processing.

Several additional directions should be pursued to improve the clinical feasibility of the PRP approach. One important issue is whether testing time can be reduced without substantially compromising reliability. Prior work on EEG to continuous speech suggests that reliability declines as the amount of data decreases (Dial & Gnanateja, 2025). In the present study, both PRP-derived measures showed moderate-to-good reliability using approximately 15 minutes of EEG data per presentation condition in each session, which is shorter than the per-condition recording durations used in some previous PRP studies (e.g., 60 minutes, Khalighinejad et al., 2017; 24 or 48 minutes, Aldag & Nogueira, 2024). This suggests that clinically practical recording times may be achievable, although the impact of further reduction in recording duration will need to be evaluated empirically. At the same time, while the ICCs we observed fell within the moderate-to-good range (0.628–0.845), some recommendations for clinical measures call for reliability of at least 0.70 (Boateng et al., 2018; Frost et al., 2007; Nunnally & Bernstein, 1994). By this standard, roughly half of the measures shown in Figure 5 would require further improvement. One possible way to increase reliability is to focus on the subset of phonemes (e.g., the Ling 6 phonemes; Glista et al., 2014) yielding the most stable PRP-derived measures and to design speech materials that maximize the occurrence of those phonemes. Finally, because PRPs have shown sensitivity to speech processing differences across listener populations, it will be important to extend this work beyond young adults with normal hearing. In the longer term, a large normative PRP database could help establish reference standards for listeners with diverse demographic and clinical characteristics.

## Conclusion

PRPs emerge as a promising new paradigm for capturing cortical speech processing under naturalistic listening conditions. In the present study, PRP-derived measures of neural differentiation among manner classes did not vary significantly across testing sessions, presentation modalities (binaural vs. monaural), or stimulus intensity levels (60 vs. 75 dB), and showed moderate-to-good reliability. In contrast, neural tracking of multiband acoustic envelope and onset features was influenced by these factors, particularly presentation modality. Presumably because PRPs reflect processing of abstract, linguistically relevant categories, measures based on PRPs show stability that makes them suitable for clinical translation, complementing conventional audiologic tests that primarily assess peripheral hearing. Future work should further improve the clinical feasibility of the PRP approach, including evaluating sparse electrode montages and shorter testing durations.

## Acknowledgments

This research was supported in part by the Pat & Shirley Ryan Family Research Acceleration Fund at Northwestern University to B.C. and J.R.M, and in part by the computational resources and staff contributions provided by the Quest high-performance computing facility at Northwestern University, which is jointly supported by the Office of the Provost, the Office of Research, and Northwestern University Information Technology. The authors thank Shengyue Xiong for assistance with data collection.

## Author Contributions

**Zhe-chen Guo**: Conceptualization; Investigation; Data curation; Formal analysis; Software; Validation; Visualization; Writing – original draft.

**Kailyn A. McFarlane**: Conceptualization; Investigation; Methodology; Data curation; Project administration; Writing – original draft.

**Jacie R. McHaney**: Conceptualization; Funding acquisition; Investigation; Methodology; Writing – original draft.

**Ishika Choksi:** Investigation; Data curation; Project administration.

**Mollee Feeney:** Investigation; Data curation; Project administration.

**Lauren Preston:** Investigation; Data curation; Project administration.

**Bharath Chandrasekaran**: Conceptualization; Funding acquisition; Project administration; Resources; Supervision; Writing – review & editing.

